# Reduced mitochondrial lipid oxidation leads to fat accumulation in myosteatosis

**DOI:** 10.1101/471979

**Authors:** Jonathan P Gumucio, Austin H Qasawa, Patrick J Ferrara, Afshan N Malik, Katsuhiko Funai, Brian McDonagh, Christopher L Mendias

## Abstract

Myosteatosis is the pathological accumulation of lipid that can occur in conjunction with atrophy and fibrosis following skeletal muscle injury. Little is known about the mechanisms by which lipid accumulates in myosteatosis, but many clinical studies have demonstrated the degree of lipid infiltration negatively correlates with muscle function and regeneration. Our objective was to determine the pathological changes that result in lipid accumulation in injured muscle fibers. We used a rat model of rotator cuff injury in this study, as the rotator cuff muscle group is particularly prone to the development of myosteatosis after injury. Muscles were collected from uninjured controls, or 10, 30, or 60 days after injury, and analyzed using a combination of muscle fiber contractility assessments, RNA sequencing, and undirected metabolomics, lipidomics and proteomics, along with bioinformatics techniques, to identify potential pathways and cellular processes that are dysregulated after rotator cuff tear. Bioinformatics analyses indicated that mitochondrial function was likely disrupted after injury. Based on these findings, and given the role that mitochondria play in lipid metabolism, we then performed targeted biochemical and imaging studies and determined that mitochondrial dysfunction and reduced fatty acid oxidation likely leads to the accumulation of lipid in myosteatosis.

## Introduction

Skeletal muscle often displays chronic degenerative changes following injury, including atrophy and weakness of muscle fibers, and fibrotic changes to the extracellular matrix (1-4). Some muscles also display a pathological accumulation of lipid in response to chronic injury or disease, which is referred to as myosteatosis or fatty degeneration (5, 6). The rotator cuff muscle group, which moves and stabilizes the shoulder joint, is particularly prone to develop myosteatosis after injury (7). Chronic tears of the rotator cuff, which nearly always involve a tear to the distal tendon, are among the most frequent upper extremity injuries in patients, with over a quarter of a million surgical repairs performed annually in the US (8, 9). Patients with these injuries have a 30% reduction in muscle fiber force production, and degenerative changes to the motor endplate consistent with partial denervation (1, 10). Lipid accumulation in injured rotator cuff muscles is correlated with muscle weakness and poor patient outcomes following surgical repair (7, 11). Additionally, for most patients the muscle does not regain strength and function despite undergoing postoperative strengthening exercises in rehabilitation, and nearly 40% of patients will actually continue to develop more atrophy and fat accumulation after the repair (7). As excess lipid appears to play a negative role in muscle regeneration, developing therapeutic interventions to reduce myosteatosis may improve outcomes for patients with chronic muscle injuries and diseases. However, the mechanisms that lead to the pathological build-up of lipid within muscle fibers are not well understood.

Our objective in the current study was to gain greater insight into the biochemical pathways and cellular factors that lead to myosteatosis after skeletal muscle injury. We used a rat model of rotator cuff tears, in which the supraspinatus muscle was avulsed from the humerus and denervated, and muscles were harvested either 10, 30 or 60 days after the injury, and compared to supraspinatus muscles of uninjured rats. We divided the study into two parts. First, we evaluated changes in muscle fiber force production and performed a broad analysis of the changes in the muscle lipidome, metabolome, transcriptome, and proteome using mass spectrometry and RNA-sequencing techniques. Bioinformatics tools were used to identify potential cellular processes and molecular targets that were disrupted as a result of rotator cuff injury. This analysis identified mitochondrial dysfunction as a strong candidate in causing pathological changes after muscle injury. Based on these results, in the second half of the study we tested the hypothesis that pathological lipid accumulation occurs in torn rotator cuff muscles due to mitochondrial dysfunction and reduced lipid oxidation.

## Methods

### Animals

This study was approved by the University of Michigan IACUC. Forty adult male Sprague Dawley rats were used in this study (N=10 per group). The rat rotator cuff tenectomy and denervation model was chosen due to anatomical similarity to humans, and because the model mimics many of the pathological changes observed in patients with chronic rotator cuff tears (1, 3, 12-14). Bilateral supraspinatus (SSP) tears were administered as previously described (12, 14, 15). Animals were deeply anesthetized with 2% isoflurane, and the surgical area was shaved and scrubbed with chlorhexidine. A deltoid-splitting transacromial approach was used to expose the supraspinatus tendon, and a tenectomy was performed to remove the tendon and prevent spontaneous reattachment to the humerus or surrounding fascia. Following tenectomy, the suprascapular nerve was located, and approximately a 3-4mm segment was removed to denervate the muscle. The deltoid muscle and skin were then closed, and the animals were allowed to recover in their cages. Rats were treated with buprenorphine twice post-operatively (0.05mg/kg, Reckitt, Parsippany, NJ, USA) and a single dose of carprofen (5mg/kg, Zoetis, Parsippany, NJ, USA). Animals were closely monitored thereafter for signs of pain or distress. After a period of either 10, 30, or 60 days, animals were anesthetized with sodium pentobarbital (40mg/kg, Vortech Pharmaceuticals, Dearborn, MI, USA), and supraspinatus muscles were harvested, weighed, and prepared for either histology, single fiber contractility, or finely minced and snap frozen in 25-50mg aliquots for biochemical and molecular biology measures. Animals were then euthanized by overdose of sodium pentobarbital followed by induction of a bilateral pneumothorax.

### Histology

Histology was performed as described (12). Distal portions of the supraspinatus were placed in tragacanth gum and snap frozen in isopentane cooled with liquid nitrogen. Ten micron sections were fixed with 4% paraformaldehyde, and then incubated in 0.2% Triton-X 100 and then stained with wheat germ agglutinin (WGA) conjugated to AlexaFluor 555 (Thermo Fisher, Waltham, MA, USA) to label the extracellular matrix, DAPI (Sigma, Saint Louis, MO, USA) to identify nuclei, and BODIPY 493/503 (Thermo Fisher) to label neutral lipids. To calculate fiber cross-sectional areas (CSAs), images of WGA and DAPI stained sections were taken on an EVOS FL microscope (Thermo Fisher), and CSAs were calculated using ImageJ (NIH). Images stained with WGA, DAPI, and BODIPY were taken using an AxioPhot system (Zeiss, Thornwood, NY, USA).

### Muscle Fiber Contractility

The contractility of chemically permeabilized muscle fibers was performed as described (1, 12). Briefly, fiber bundles were dissected from the proximal region of SSP muscles, placed in skinning solution for 30 min, and then in storage solution for 16 h at 4°C, followed by storage at −80°C. On the day of contractility testing, samples were thawed slowly on ice, and individual fibers were pulled from bundles using fine mirror-finished forceps. Fibers were then placed in a chamber containing relaxing solution and secured at one end to a servomotor (Aurora Scientific, Aurora, ON, Canada) and the other end to a force transducer (Aurora Scientific) with two ties of 10-0 monofilament nylon suture. Using a laser diffraction measurement system, fiber length was adjusted to obtain a sarcomere length of 2.5μm. The average fiber CSA was calculated assuming an elliptical cross-section, with diameters obtained at five positions along the length of the fiber from high-magnification images taken from top and side views. Maximum isometric force (Fo) was elicited by immersing the fiber in a high calcium activation solution. Specific maximum isometric force (sFo) was calculated by dividing Fo by fiber CSA. Fibers were categorized as fast or slow by examining their force response to rapid, constant-velocity shortening, and ten to twenty fast fibers were tested from each SSP muscle.

### Western blots

Western blots were performed as described (16). Protein was isolated from snap frozen 25mg aliquots of muscle tissue and placed in 500μL of a solution containing TPER (Thermo Fisher) with 1% NP-40 and 1% protease-phosphatase inhibitor (Thermo Fisher). Samples were homogenized, vortexed at 4°C for 30 min, then centrifuged at 12,000g for 15 min at 4°C. The supernatant was collected and saved at −80°C until use. The pellets of insoluble proteins were saved for hydroxyproline assays, as described below. Protein concentration was determined with a BCA assay (Thermo Fisher). Twenty micrograms of protein were loaded into 6-12% polyacrylamide gels, and subjected to electrophoretic separation. Proteins were transferred onto nitrocellulose membranes (Bio-Rad) and blocked for 1h in 3-5% skim milk. Membranes were rinsed and incubated overnight in primary antibodies at a concentration of 1:1000, against either glutathione (Virogen, Watertown, MA, USA, 101A) succinate dehydrogenase-A (SDHA, Cell Signaling, Danvers, MA, USA, 11998), cytochrome c oxidase subunit 4 (COX4, Cell Signaling 4850), phospho-insulin-like growth factor-1 receptor (p-IGF1R) Y^1135^ (Cell Signaling 3918), IGF1R (Cell Signaling 9750), phospho-ERK1/2 T^202^/Y^204^ (Cell Signaling 4370), ERK1/2 (Cell Signaling 4695), phospho-Akt S^473^ (Cell Signaling 4060), Akt (Cell Signaling 4691), phospho-p70S6 Kinase (p-p70S6K) T^389^ (Cell Signaling 9234), phospho-p70^S6K^ T^421^/S^424^ (Cell Signaling 9204), p70^S6K^ (Cell Signaling 2708), phospho-ribosomal protein S6 (p-rpS6) S^235^/S^236^ (Cell Signaling 4858), rpS6 (Cell Signaling 2217), p62 (AbCam, Cambridge, UK, 129012), ULK1 (AbCam 128859), PINK1 (AbCam 23707), Parkin (AbCam 15954), Prdx3 (AbCam 128953), or Prdx6 (AbCam 133348). After primary antibody incubation, membranes were rinsed and incubated in goat anti-rabbit or goat anti-mouse horseradish peroxidase-conjugated secondary antibodies (from either AbCam or Cell Signaling). Proteins were detected using Clarity Western ECL Substrate (Bio-Rad) or Super Signal West Dura (Thermo Fisher), and imaged and quantified using a ChemiDoc chemiluminescent detection system (Bio-Rad). Coomassie or Ponceau S staining of membranes was performed to verify equal protein loading.

### Hydroxyproline content

Hydroxyproline was measured from the insoluble fraction remaining from the isolation of soluble proteins for western blots. Pellets were dried at 100°C overnight, weighed immediately, and then hydrolyzed in 12M hydrochloric acid. Samples were then evaporated in a SpeedVac (Thermo Fisher) and hydroxyproline was determined using a colorimetric assay as described previously (17).

### Electron Microscopy

Portions of distal supraspinatus muscles were fixed in 1% tannic acid, 1% glutaraldehyde in Sorenson’s buffer, followed by post-fixation in 2% osmium tetroxide (Electron Microscopy Sciences, Hatfield, PA, USA). Samples were then dehydrated using a graded ethanol series and embedded in EMBed 812 (Electron Microscopy Sciences) using a graded resin and propylene oxide series. One micron transverse sections were cut with a diamond knife ultramicrotome, and imaged using a JEOL 1400-plus transmission electron microscope (JEOL USA, Peabody, MA, USA) with a high-resolution AMT digital camera (AMT, Woburn, MA, USA).

### Lipidomics and metabolomics

The University of Michigan Metabolomics Core performed mass spectrometry based shotgun lipidomics and metabolomics measurements from snap frozen, homogenized muscle samples as described (18, 19). For lipidomics, lipids were extracted from samples with a solvent mixture consisting of 2:2:2 (v/v/v) methanol:dichloromethane:water mixture at room temperature after adding internal standard mixture. After drying, the samples were resuspended in a solution containing 1:5:85 (v/v/v) acetonitrile:water:isopropanol and 10mM ammonium acetate. Samples were then subjected to liquid chromatrography-mass spectrometry (LC-MS), and MS peaks were matched in-silico with LipidBlast (20). Quantification was performed by Multiquant software (AB-SCIEX, Framingham, MA, USA). For metabolomics, metabolites were extracted from frozen muscle in a solvent mixture containing 8:1:1 methanol:chloroform:water (v/v/v). Reverse phase liquid chromatography-tandem quadrupole MS was used to measure acylcarnitines (21). Other metabolites were derivatized and analyzed with gas chromatography-MS. Quantification of metabolites was performed using Masshunter Quantitative Analysis software (Agilent Technologies, Santa Clara, CA, USA). Normalized abundance data for all measured metabolites is provided in Supplemental Material 1.

### RNA Sequencing (RNA-seq) and Gene Expression

Snap frozen aliquots of muscle tissue were homogenized in QIAzol (Qiagen, Valencia, CA) and isolated using a miRNeasy kit (Qiagen). RNA concentration was determined using a NanoDrop 2000 (Thermo Fisher). For each sample, 250ng total RNA was delivered to the University of Michigan Sequencing Core for RNA sequencing (RNA-seq) analysis. Sample concentrations were normalized and cDNA pools were created for each sample, and then subsequently tagged with a barcoded oligo adapter to allow for sample specific resolution. Sequencing was carried out using an Illumina HiSeq 2500 platform (Illumina, San Diego, CA, USA) with 50bp single end reads. Raw RNA-seq data was quality checked using FastQC v0.10.0 (Barbraham Bioinformatics, Cambridge, UK). Alignment to the reference genome (rn5, UCSC), differential expression based on counts per million mapped reads (CPM), and post-analysis diagnostics were carried out using the Tuxedo Suite software package (22). RNA-seq data has been deposited to NIH GEO (accession GSE103266), and normalized CPM values for all measured genes is provided in Supplemental Material 1.

To validate fold-change RNA-seq data, we performed quantitative PCR (qPCR) on a select set of genes. RNA was reverse transcribed into cDNA with iScript Reverse Transcription Supermix (Bio-Rad, Hercules, CA, USA). Amplification of cDNA was performed in a CFX96 real-time thermal cycler (Bio-Rad) using iTaq Universal SYBR Green Supermix (Bio-Rad). Target gene expression was normalized to the stable housekeeping gene Eif2b2, and further normalized to 0d samples using the 2^-AACt^ method. Primer sequences are provided in Supplemental Material 2.

### Proteomics

Label-free proteomic analysis was performed as described (23). Aliquots were homogenized in 50mM ammonium bicarbonate containing 25mM N-ethylmaleimide (d(0) NEM), pH 8. Protein lysates were prepared by centrifugation at 15,000g for 10min at 4°C and protein concentrations were calculated using a Bradford assay (BioRad) with BSA as a standard. Excess d(0) NEM was removed using Zeba desalting columns (Thermo Scientific), and protein concentrations were determined again by Bradford assay as before. A total of 100μg of protein extract was diluted to 160μl with 25mM ammonium bicarbonate and denatured by the addition of 10μl of 1% RapiGest (Waters, Manchester, UK) in 25 mM ammonium bicarbonate and incubated at 80°C for 10min with shaking. Then 10μl of a 100mM solution of Tris(2-carboxyethyl)phosphine hydrochloride (TCEP) was added to reduce reversibly oxidised Cys residues, followed by incubation at 60°C for 10min. Newly reduced Cys were then alkylated by addition of d(5) NEM and incubated at room temperature for 30min. An aliquot of the samples was used at this point to check procedure by SDS-PAGE. Proteolytic digestion was performed by addition of trypsin followed by overnight incubation at 37°C. Digestion was terminated and RapiGest removed by acidification (3μ1 of TFA and incubated at 37°C for 45min) and centrifugation (15,000g for 15min).

Samples were analyzed using an Ultimate 3000 RSLC Nano system (Thermo Scientific) coupled to a QExactive mass spectrometer (Thermo Scientific). A total of 2μl of sample corresponding to 1 μg of protein was diluted in 18μl buffer (97% H2O, 3% MeCN and 0.1 % formic acid v/v) and 5μ1 (250ng of protein) was loaded onto the trapping column (PepMap 100, C18, 75μm x 20mm, Thermo Fisher) using partial loop injection for 7min at a flow rate of 4μL/min with 0.1% (v/v) TFA. Samples were resolved on the analytical column (EASY-Spray C18 75μm x 400mm, 2μm column, Thermo Fisher) using gradient of 97% A (0.1% formic acid) and 3% B (99.9% ACN and 0.1% formic acid) to 60% A and 40% B over 120 min at a flow rate of 300nL/min. Data dependent acquisition consisted of a 70,000 resolution full-scan MS scan (AGC set to 10^6^ ions with a maximum fill time of 250ms) and the 10 most abundant peaks were selected for MS/MS using a 17,000 resolution scan (AGC set to 5 × 10^4^ ions with a maximum fill time of 250ms) with an ion selection window of 3*m/z* and normalized collision energy of 30. Repeated selection of peptides for MS/MS was avoided by a 30sec dynamic exclusion window.

Label-free relative quantification was performed using PEAKS7 software (Bioinformatics Solutions Inc., Waterloo, Ontario, Canada). The acquired Thermo RAW data files were searched against UniProt rat database (2015-29-07, 32,991 sequences) and analyzed using the following parameters: peptide mass tolerance 10ppm; fragment mass tolerance 0.01Da, 1+, 2+, 3= ions; missed cleavages. Variable modifications included in search were: d(0) NEM, d(5) NEM, mon-, di-, trioxidation of Cys residues and oxidation of methionine with a FDR of < 1% and searched against the UniProt rat database. Normalization was carried out using the total ion current. PEAKS7 software includes a post-translational modification (PTM) algorithm applying the de novo sequencing module to search for a limited number of PTMs. All identified PTMs using this method adhere to the above search criteria and FDR validation. All proteomics data has been deposited to the Proteome Xchange Consortium (accession PXD009034), and normalized abundance data for all measured proteins is provided in Supplemental Material 1.

### Bioinformatics

Expression data from RNA-seq measurements was imported into Ingenuity Pathway Analysis (IPA) software (Qiagen) to assist in predicting cellular and molecular pathways and processes involved in myosteatosis. For metabolomics, lipidomics, and proteomics measures, MS peak data was imported into MetaboAnalyst 4.0 software (24) for data visualization, principal component analysis (PCA), and statistical analyses.

### Mitochondrial DNA (mtDNA) measurements

Mitochondrial genome copy number quantification was performed with modifications from a previous study (25). Total DNA was isolated from aliquots of muscle tissue using the DNeasy Blood and Tissue Kit (Qiagen). DNA was amplified via qPCR as described above with custom primers for mtDNA and beta-2-microglobulin (B2M) genomic DNA (Supplemental Material 2). Using known standards of rat DNA, standard curves were generated for each primer set to determine number of copies of either mtDNA or gDNA. The ratio of mtDNA to gDNA was calculated per sample and averaged across groups.

### Mitochondrial enzymatic assays

Fifty milligrams of freshly minced supraspinatus tissue was isolated and placed in 500μL of ice cold PBS containing protease inhibitor (Thermo Fisher). Samples were homogenized and then 100μL of 10% laurel maltoside, and 400μL cold PBS with protease inhibitor was further added. Samples were incubated on ice for 30 min and then centrifuged at 12,000g for 20 min at 4°C. Samples were stored at −80°C until use. On the day of measurement, samples were thawed on ice, and protein concentration was determined using a BCA assay (Thermo Fisher). Colorimetric enzymatic assays for mitochondrial complex I, II, and IV were performed as per the protocol acquired from the manufacturer (AbCam). Sample concentration was optimized per assay in pilot experiments, and equal protein was loaded per well. Samples were measured in duplicate in a BioTek Epoch2 microplate spectrophotometer (BioTek, Winooski, VT, USA). Optical density of enzymatic activity per minute (mOD/min) was calculated per the manufacturer protocol.

### Pyruvate and palmitate oxidation measurements

Samples were placed into ice-cold medium containing 250mM sucrose, 1mM EDTA, 10mM Tris-HCl, and 2mM ATP (pH 7.4). To isolate intact mitochondria, muscles were finely minced in 20μL of medium per milligram of muscle, and homogenized in a glass homogenization tube with a motor-driven Teflon pestle (26). Pyruvate and palmitate oxidation assays were performed as previously described with minor modification (27). Briefly, 80μL of tissue homogenate was added to incubation wells on a sealed, modified, 48-well plate with a channel cut between the adjacent trap wells. The trap wells contained 200μL of 1M sodium hydroxide for the collection of liberated ^14^CO_2_. Two volumes of incubation buffer (100mM sucrose, 10mM Tris-HCl, 5mM potassium phosphate, 80mM potassium chloride, 1mM magnesium chloride, 0.1mM malate, 2mM ATP, 1mM dithiothreitol, 0.2mM EDTA, 1mM L-carnitine, 0.05mM coenzyme A, and 0.5% fatty acid-free bovine serum albumin, pH 7.4) was added to the wells to initiate the reaction with 1mM pyruvate ([2-^14^C]pyruvate at 0.5μCi/mL) or 0.2mM palmitate ([1-^14^C]palmitate at 0.5μCi/mL). Following 60 min of incubation at 37°C, 100μl of 70% perchloric acid was added to terminate the reaction. The trap wells were sampled for label incorporation into ^14^CO_2_, which was determined by scintillation counting using 4mL of Uniscint BD (National Diagnostics, Atlanta, GA, USA).

### Statistics

Differences between groups were measured using one-way ANOVAs (α=0.05) followed by Fisher’s LSD post hoc sorting (α=0.05), or Kruskal-Wallis tests (α=0.05) followed by Dunn’s post-hoc sorting (α=0.05). A FDR multiple observation adjustment (q<0.05) was applied to metabolomic, lipidomic, proteomic, and RNA-seq data. Statistical analyses were performed using Prism 6.0 (GraphPad), Tuxedo, or MetaboAnalyst 4.0 (24).

## Results

Following muscle injury, the mass of muscles and size of muscle fibers decreased over time (Figure 1A-B). Collagen content also accumulated after muscle injury, with a near 4-fold increase at 30 and 60 days compared to controls (Figure 1C). An accumulation of lipids within muscle fibers was noted after muscle injury, both on histology and electron microscopy, as well as disrupted myofibril architecture (Figure 1D and 2A). Centrally located nuclei were also noted 10 days after injury (Figure 1D). In addition to a reduction in size, muscle fibers were also weaker, both in terms of absolute maximum isometric force and specific force (Figure 2B-D).

**Figure 1.**
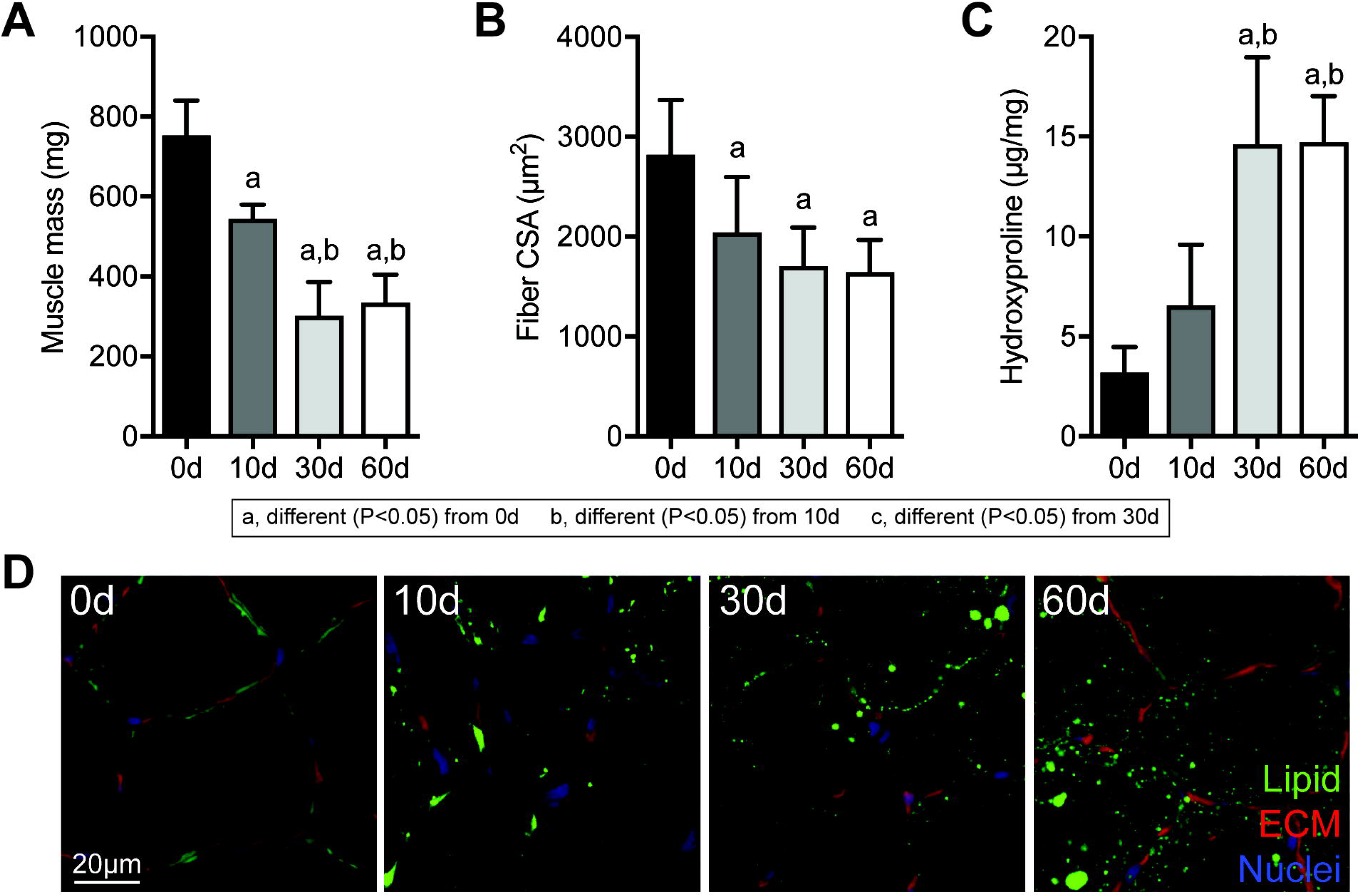
Changes in muscle mass, fiber size, collagen content, and lipid accumulation after rotator cuff tear. (A) Mass, (B) muscle fiber histology cross-sectional area (CSA) values, and (C) hydroxyproline content of muscles following rotator cuff tear. (D) Representative histology of areas of muscle demonstrating lipid accumulation, with neutral lipids in green, extracellular matrix in red, and nuclei in blue. Data are presented mean+SD, N>6 muscles per group. Post-hoc sorting (P<0.05): a, different from 0d; b, different from 10d; c, different from 30d.

**Figure 2.**
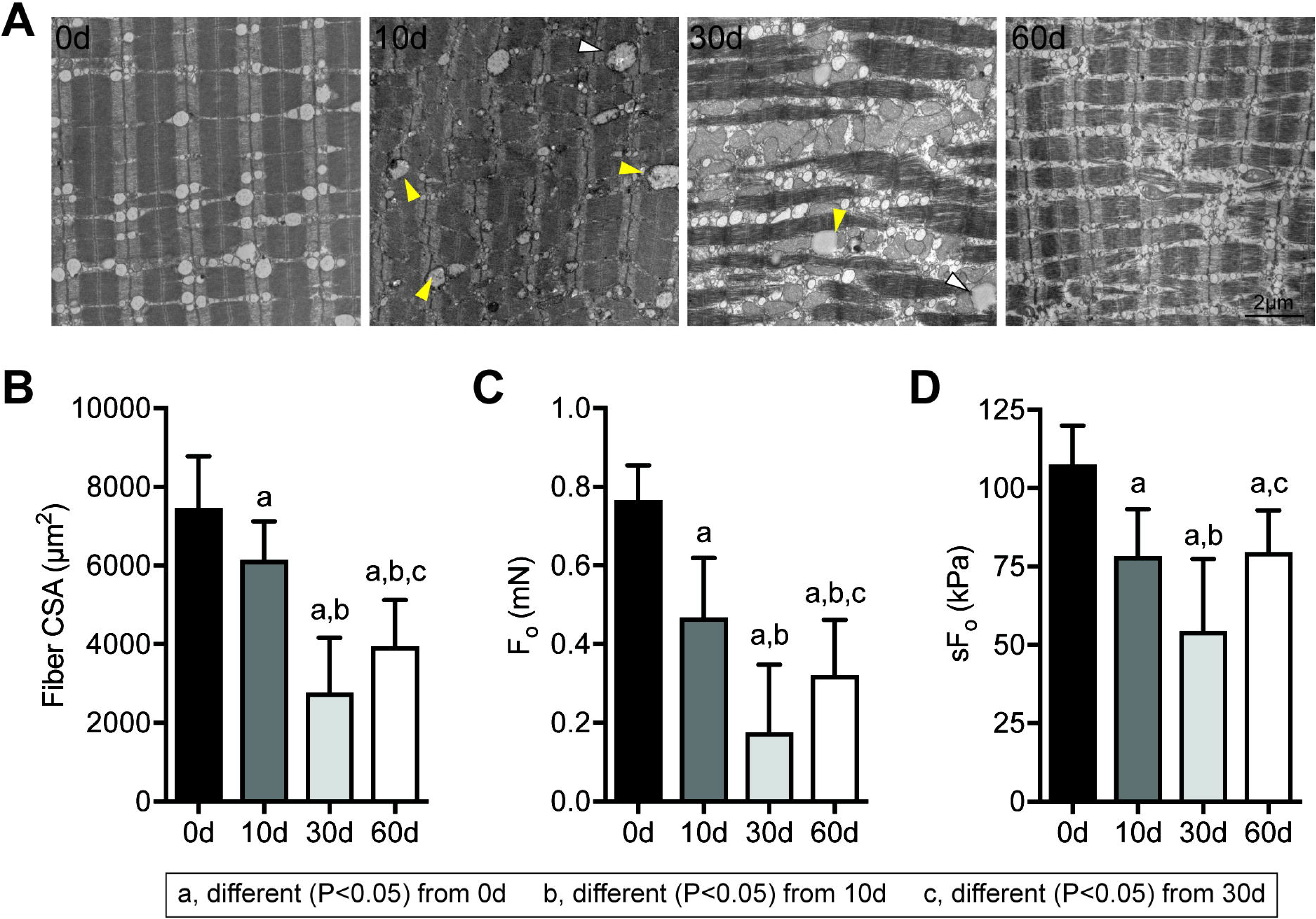
Changes in myofibrillar architecture and muscle fiber contractility after rotator cuff tear. (A) Representative electron micrographs taken in the middle of the fiber demonstrating disrupted sarcomere ultrastructure in supraspinatus muscles following injury. Arrowheads indicate lipid-laden mitochondria. (B) Cross-sectional area (CSA), (C) maximum isometric force (Fo), and (D) specific force (sFo) of permeabilized muscle fibers from control and injured muscles. N=10 muscles per group. Post-hoc sorting (P<0.05): a, different from 0d; b, different from 10d; c, different from 30d.

Given the reduction in muscle fiber size, we then measured activation of the IGF1 and ERK pathways, due to their importance in maintaining muscle mass (Figure 3A-G). Reduced proximal IGF1 and ERK signaling was observed after muscle injury, with corresponding reductions in Akt S^473^ and p70S6K T^421^/S^424^, although rpS6 S^235^/S^236^ phosphorylation was increased 10 and 30 days after muscle injury (Figure 3A-G).

**Figure 3.**
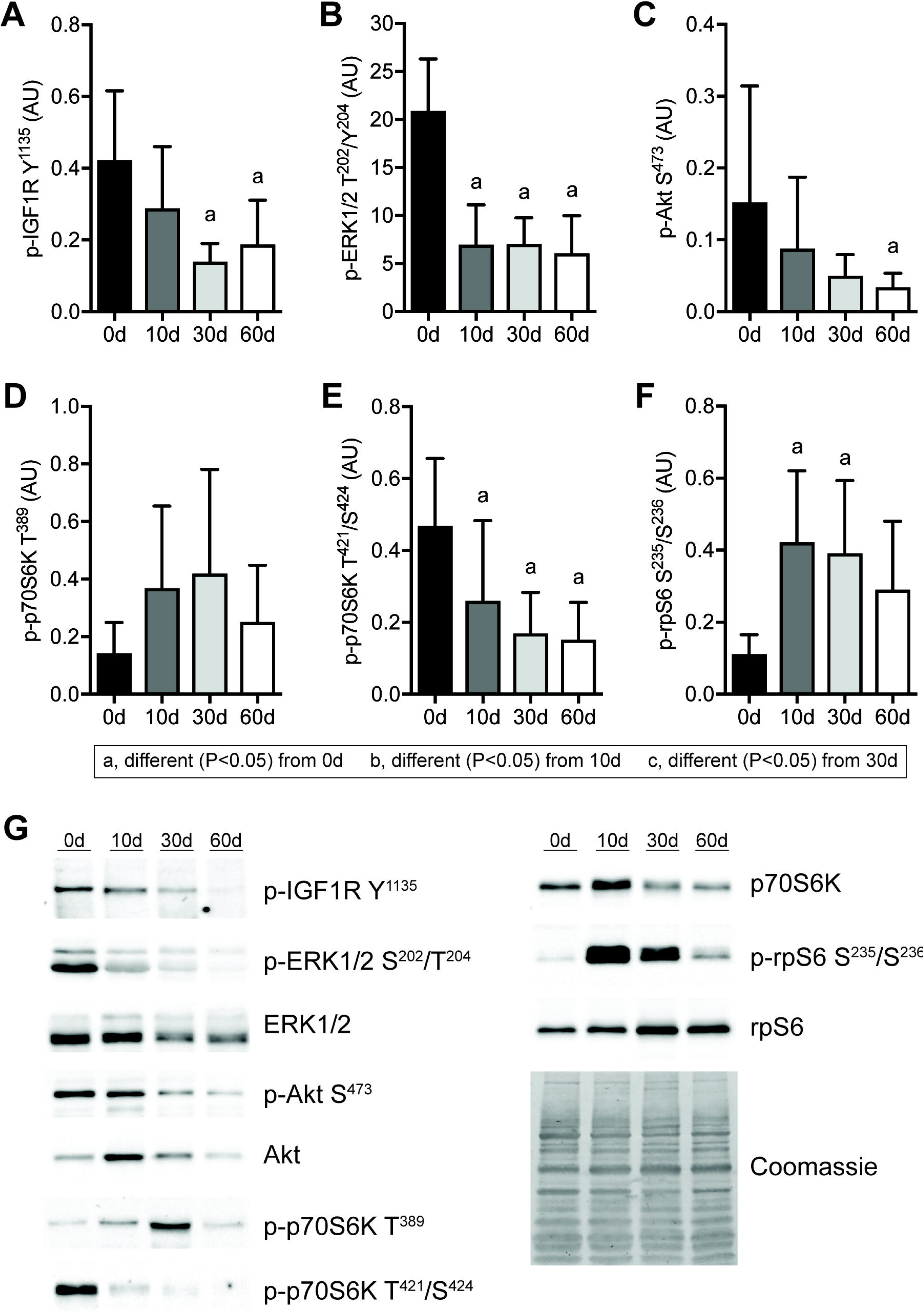
Changes in IGF1 signaling after rotator cuff tear. Quantification of band densitometry of (A) p-IGF1R Y^1135^, (B) p-ERK1/2 T^202^/Y^204^, (C) p-Akt S^473^, (D) p-p70S6K T^389^, (E) p-p70S6K T^421^/S^424^, (F) p-rpS6 S^235^/S^236^, normalized to total protein, following rotator cuff tear. (G) Representative phospho and total protein Western blots, with a Coomassie stained membrane as a loading control. Data are presented mean+SD, N=6 per group. Post-hoc sorting (P<0.05): a, different from 0d; b, different from 10d; c, different from 30d.

To further quantify and explore the grossly apparent lipid that accumulated after rotator cuff tear, we performed shotgun lipidomics and detected 457 lipid species, with summary data presented in Figure 4. In general, PCA analysis demonstrated that the 10 day injury group diverged somewhat from uninjured muscles, while the 30 and 60 day injury groups were fairly distinct from uninjured muscles but had some overlap with the 10 day group (Figure 4A). For free fatty acids (FFAs), there was an increase in oleate, palmitate, and stearate 30 and 60 days after rotator cuff tear (Figure 4B). Triglycerides were the major class of lipid species measured in muscle tissues and increased by over 3-fold at the 60 day time point (Figure 4C). Ceramide phosphates, sphingomyelins, phosphatidylglycerols, phosphatidylinositols, phosphatidylserines (Figure 4G-H, J-L) displayed similar patterns of change as triglycerides, while diglycerides, monoglycerides, cholesterol esters, phosphatidic acids, phosphatidylcholines, phosphatidylethanolamines, lysophosphatidylcholines, lysophosphatidylethanolamines, and plasmenyl-phosphatidylcholines (Figure 4D-F, I, M-Q) displayed biphasic responses. No difference in plasmenyl-phosphatidylethanolamines (Figure 4R) were observed.

**Figure 4.**
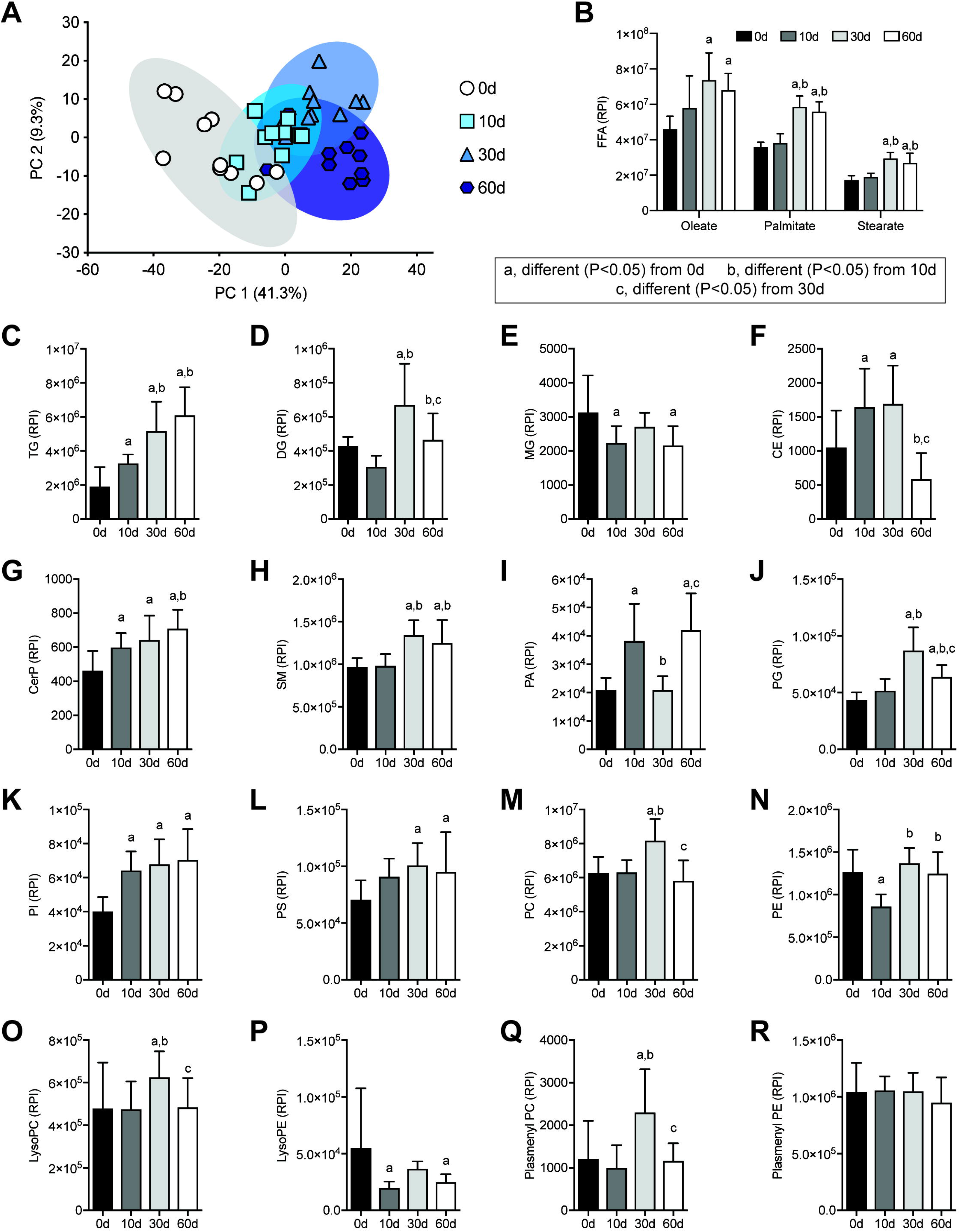
Changes in lipid species after rotator cuff tear. (A) Principal component analysis of groups C-R. Levels of (B) free fatty acids (FFA), (C) triglycerides (TG), (D) diglycerides (DG), (E) monoglycerides (MG), (F) cholesterol ester (CE), (G) ceramide phosphates (CerP), (H) sphingomyelins (SM), (I) phosphatidic acids (PA), (J) phosphatidylglycerols (PG), (K) phosphatidylinositols (PI), (L) phosphatidylserines (PS), (M) phosphatidylcholines (PC), (N) phosphatidylethanolamines (PE), (O) lysophosphatidylcholines (LysoPC), (P) lysophosphatidylethanolamines (LysoPE), (Q) plasmenyl phosphatidylcholines (Plasmenyl PC), and (R) plasmenyl phosphatidylethanolamines (Plasmenyl PE), as measured by mass spectrometry and presented as relative peak intensity (RPI). Data are presented mean+SD, N=10 muscles per group for A and C-R, and N=5 muscles per group for B. Post-hoc sorting (P<0.05): a, different from 0d; b, different from 10d; c, different from 30d.

Next, we analyzed the levels of 68 small molecule metabolites important for muscle function. Select metabolites are presented in Figure 5, with additional species listed in Supplemental Material 1. There was a general similarity of metabolites at the 0 and 10 day time points, and at the 30 and 60 day time points (Figure 5A). Several nucleoside and nucleotide metabolites, such as ADP, CMP, GMP, NAD, and UMP decreased after rotator cuff tear (Figure 5B). An increase in metabolites involved in glycolysis and pentose phosphate metabolism, such as fructose-bisphosphate, hexoses, hexose-phosphate, and phosphoenolpyruvate increased at 30 and 60 days after muscle injury (Figure 5D). There were changes in only a few free amino acids such as alanine, aspartate, threonine, and tryptophan (Figure 5E), and no changes were observed in the creatine phosphate shuttle, Kreb’s cycle metabolites, or free glutathione after injury (Figure 5C, F, and H).

**Figure 5.**
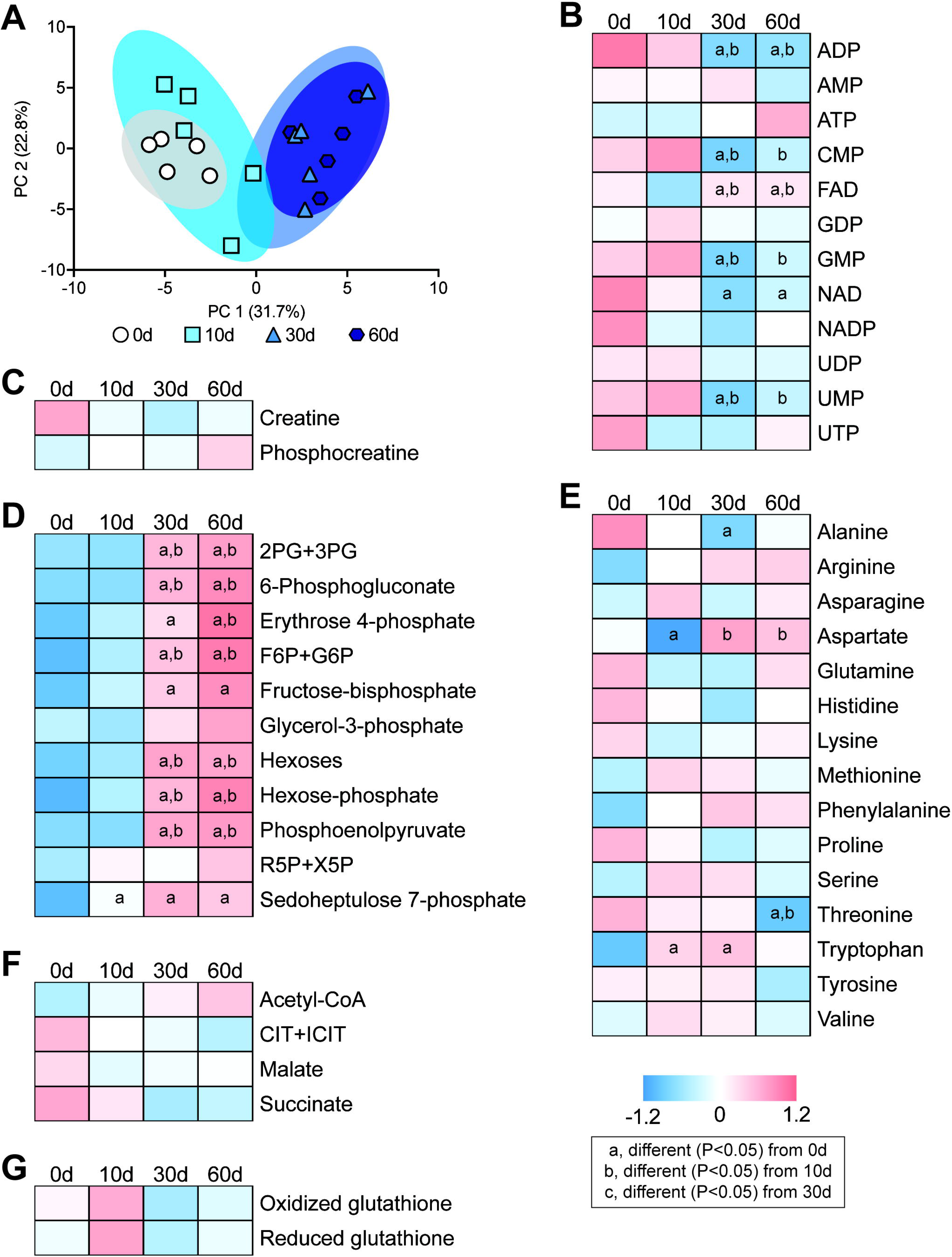
Changes in metabolites after rotator cuff tear. (A) Principal component analysis of groups. Baseline normalized heatmaps demonstrating levels of selected (B) nucleotide and nucleoside metabolites, (C) creatine and phosphocreatine, (D) glycolysis and pentose phosphate metabolites, (E) free amino acids, (F) Kreb’s cycle metabolites, and (G) oxidized and reduced glutathione, as measured by mass spectrometry. N=5 muscles per group. Post-hoc sorting (P<0.05): a, different from 0d; b, different from 10d; c, different from 30d.

We then performed RNA-seq to analyze global changes in the transcriptome of injured muscles. The PCA analyses demonstrated divergence across the different time points, with the 10 and 30 day time points demonstrating the most difference from controls (Figure 6A). IPA was used to identify potential biological pathways and functions that were affected by rotator cuff tear, and we observed several processes related to lipid metabolism and mitochondrial function, reactive oxygen species (ROS) production, glycolysis, muscle contraction, extracellular matrix (ECM) production, and inflammation that were predicted to be differentially regulated in myosteatosis (Figure 6B). We then selected numerous genes related to these processes from RNA-seq to further explore, and report in Figure 6. We also performed qPCR to validate selected genes from the different categories, and generally observed similar trends in differential regulation between RNA-seq and qPCR data (Table 1).

**Table 1.**
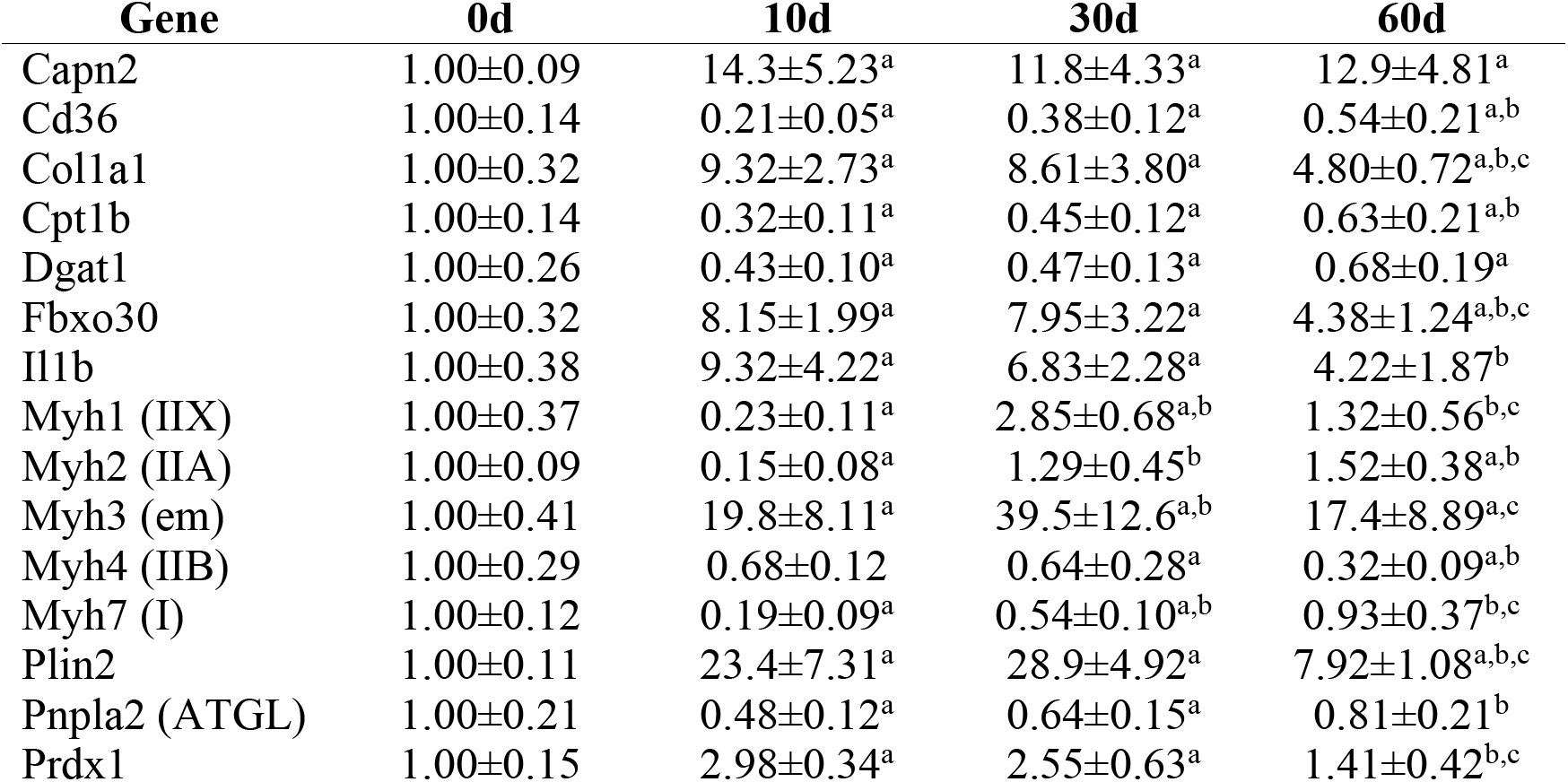
Changes in gene expression measured by qPCR. Gene expression is normalized to the stable housekeeping gene Eif2b2, and further normalized to the 0d group. Values are mean±CV, N=4 per group. Post-hoc sorting (P<0.05): a, different from 0d; b, different from 10d; c, different from 30d.

**Figure 6.**
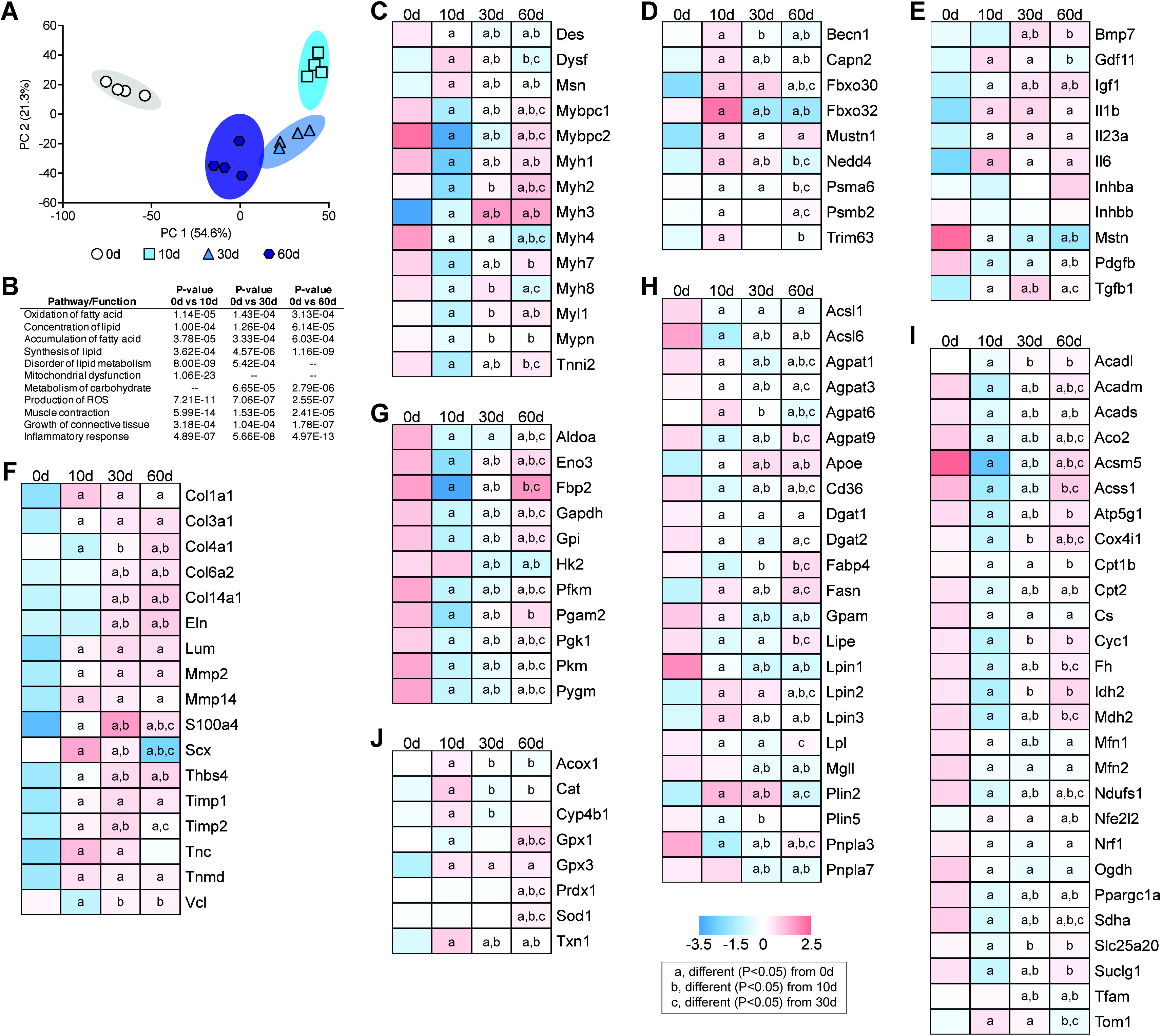
Changes in the transcriptome after rotator cuff tear. (A) Principal component analysis of overall changes across the transcriptome. (B) P-values of selected biological pathways and functions identified from Ingenuity Pathway Analysis. Levels of selected (C) contractile and structural genes, (D) autophagy and atrophy genes, (E) growth factors and cytokines, (F) extracellular matrix genes, (G) glycolysis genes, (H) lipid storage and mobilization genes, (I) mitochondrial and oxidative metabolism genes, (J) reactive oxygen species and peroxisomal oxidation genes, as measured by RNA-seq. N=4 muscles per group. Post-hoc sorting (P<0.05) a, different from 0d; b, different from 10d; c, different from 30d.

Ten days after rotator cuff tear, there was a general trend for a downregulation in genes involved in muscle contraction, such as the myosin heavy chain genes (*Myh1, Myh2, Myh4, and Myh7*) and other sarcomeric genes (*Mybpc1, Mybpc2, Myl1, Mypn, Tnni2*), as well as an increase in the membrane repair gene dysferlin (*Dysf*, Figure 6C and Table 1). Many of these genes returned to baseline by 60 days, although of note embryonic myosin heavy chain (*Myh3*) remained elevated throughout the study (Figure 6C and Table 1).

Numerous autophagy and proteosome genes such as beclin-1 (*Becn1*), m-calpain (*Capn2*), and MUSA-1 (*Fbxo30*), were induced 10 days after injury (Figure 6D), as were several genes involved with inflammation, atrophy, and fibrosis, like GDF11, ILip, and TGFp (Figure 6D-E). However, no difference in the atrophy-inducing signaling molecules activin A (*Inhba*) or B (*Inhbb*) and a downregulation in myostatin (MSTN) was observed. IGF1 and BMP7, which can activate pathways that promote muscle hypertrophy, were upregulated after rotator cuff tear (Figure 6E).

Many extracellular matrix genes, such as types I, III, IV, VI, and XIV collagen, elastin (*Eln*), lumican (*Lum*), thrombospondin 4 (*Thsbs4*), and tenascin C were upregulated at the 10 or 30 day time points, as were the fibroblast markers S100a4 and scleraxis (Figure 6F).

Genes involved in glycolysis, such as phosphofructokinase (*Pfkm*), aldolase A (*Aldoa*), GAPDH, phosphoglycerate (*Pgk1*), and enolase 3 (*Eno3*), were downregulated at 10 days, with some recovery through to 60 days after injury (Figure 6G).

For fatty acid uptake and triglyceride synthesis genes, there was a general trend for downregulation after rotator cuff tear. Lipoprotein lipase, which hydrolyzes extracellular triglycerides into fatty acids, and the fatty acid transporter CD36 was downregulated after injury (Figure 6H). Acyl-CoA synthetases (*Acsl*) 1 and 6, which are the major enzymes that conjugate fatty acids into fatty acyl Co-As in skeletal muscle were downregulated at all time points after injury, as were AGPAT9/GPAT3 and GPAM, which convert fatty acyl Co-As into lysophosphatidic acids (Figure 6H). Additional genes that were downregulated include the AGPAT1, AGPAT3, and AGPAT6 enzymes that produce phosphatidic acid from lysophosphatidic acid, lipin 1-3 which dephosphorylate phosphatidic acid to produce diglyceride, and DGAT1 and DGAT2 that produce TG from DG (Figure 6H). While TG synthesis genes were downregulated, the lipid droplet coating gene perilipin 2 (*Plin2*) was upregulated 10 days after rotator cuff tear, while perilipin 5 (*Plin5*) was downregulated. For genes involved with hydrolyzing lipids stored in droplets into fatty acids, including the patatin domain containing proteins (*Plnla2*, *Pnpla3, Pnpla7*), hormone sensitive lipase (*Lipe*), and monoglyceride lipase (*Mgll)*, were all downregulated after injury (Figure 6J and Table 1).

Enzymes that are responsible for the transport of fatty acids into the mitochondria, and their conversion into fatty acyl-CoAs, ACSM5, ACSS1, CPT1b, carnitine acylcarnitine translocase (*Slc25a20*), and CPT2 were downregulated at 10 and 30 days after injury, although some recovery occurred by 60 days (Figure 6I). For genes involved in β-oxidation of fatty acyl Co-As within the mitochondria, the short, medium and long-chain acyl Co-A dehydrogenases (*Acads*, *Acadm*, and *Acadl*) were all downregulated at 10 and 30 days after injury (Figure 6I).

Similar to β-oxidation genes, Kreb’s Cycle and oxidative phosphorylation transcripts were also downregulated after muscle injury, including citrate synthase (*Cs*), aconitase (*Aco2*), isocitrate dehydrogenase (*Idh2*), 2-oxoglutarate dehydrogenase (*Ogdh*), succinyl CoA synthetase (*Suclg1*), succinate dehydrogenase (*Sdha*), fumarase (*Fh*), malate dehydrogenase (*Mdh2*), NDUFS1, cytochrome C1 (*Cyc1*), COX4 (*Cox4i1*), and ATP5G1 (Figure 6I). Genes involved in mitochondrial biogenesis, such as PGC1a (*Ppargc1a*) and NRF1 were downregulated 10 days after injury, with no difference in TFAM expression was observed until 30 and 60 days (Figure 6I). MFN1 and MFN2, which promote mitochondrial fission, were also downregulated after rotator cuff tear (Figure 6I).

Genes involved in ROS metabolism were generally upregulated 10 days after injury, including catalase (*Cat*), glutathione peroxidase 3 (*Gpx3*), and thioredoxin (*Txn*), while others such as glutathione peroxidase 1 (*Gpx1*), peroxiredoxin 1 (*Prdx1*), and superoxide dismutase (*Sod1*), were elevated at 60 days compared to controls (Figure 6J). Although mitochondrial P-oxidation genes were downregulated, oxidation of fatty acids can also occur in peroxisomes, in a process mediated by ACOX1, which was upregulated 10 days after injury. In addition to initiating the oxidation of fatty acyl Co-As, ACOX1 produces H2O2 that is buffered in peroxisomes by catalase, which was markedly upregulated 10 days after injury (Figure 6J). ω-oxidation of fatty acids can also occur as a way to generate acetyl Co-A during times of stress, in a process regulated by CYP4B1, which was upregulated after injury (Figure 6J).

Following the analysis of transcriptional changes, we next sought to determine changes in the proteome after muscle injury. A selection of the 632 proteins detected are presented in Figure 7. The PCA analysis demonstrated that the control and 10 day groups were unique from each other and the 30 and 60 day groups, while the 30 and 60 day groups showed some overlap (Figure 7A). In most cases, we saw general agreement between observed changes in mRNAs and the proteins they encode, including those involved with protein synthesis and atrophy (Figure 7B), contractile and structural proteins (Figure 7C), extracellular matrix proteins (Figure 7D), metabolic proteins (Figure 7E), and reactive oxygen species proteins (Figure 7F).

**Figure 7.**
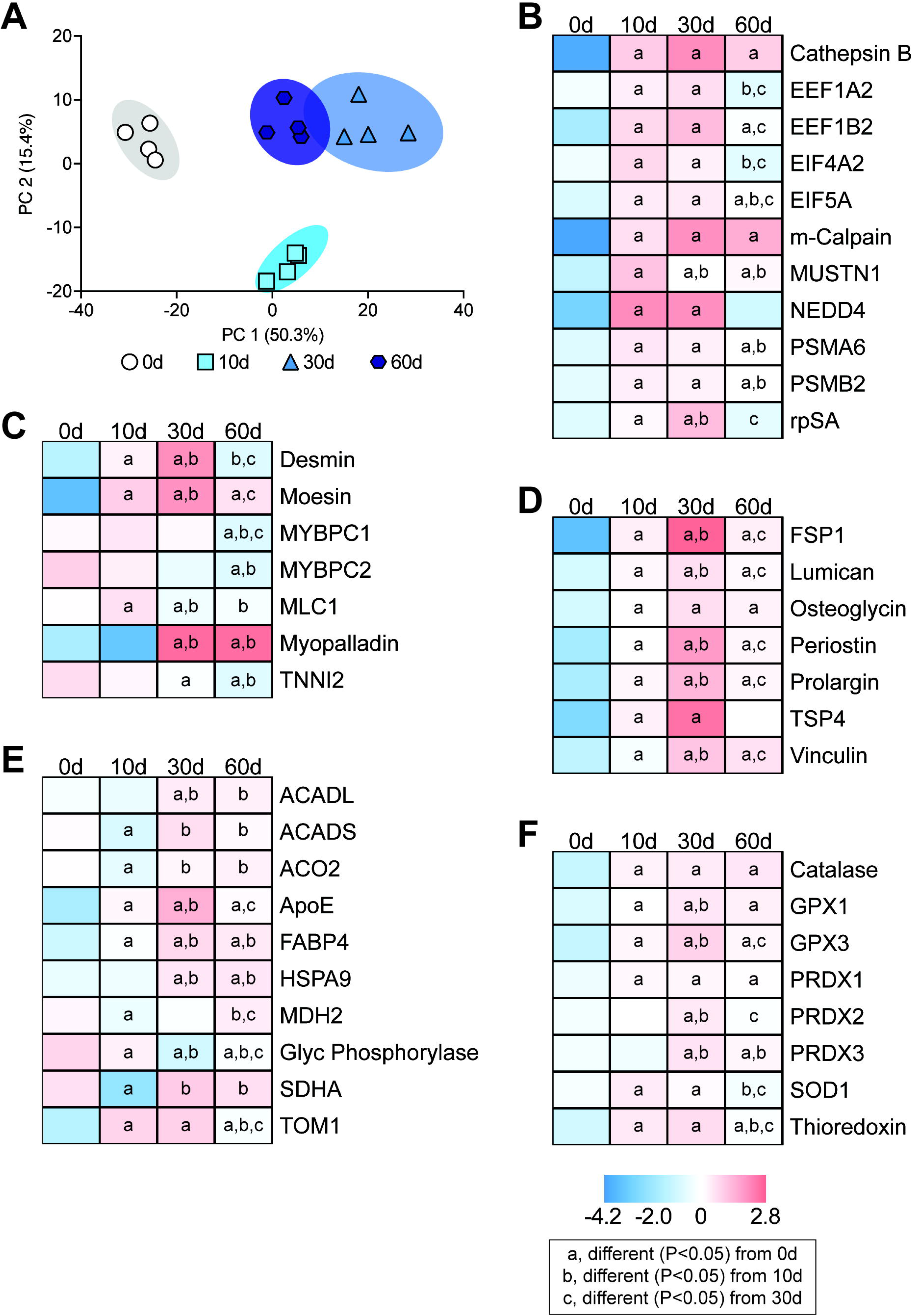
Changes in the proteome after rotator cuff tear. (A) Principal component analysis of overall changes across the proteome. Levels of selected (B) protein synthesis and atrophy proteins, (C) contractile and structural proteins, (D) extracellular matrix proteins, (E) metabolic proteins, and (F) reactive oxygen species proteins, as measured by mass spectrometry. N=4 muscles per group. Post-hoc sorting (P<d0.05): a, different from 0d; b, different from 10d; c, different from 30d.

Finally, as metabolomic, transcriptomic, and proteomic analyses suggested alterations in lipid oxidation, we sought to evaluate the abundance of lipid species and proteins that are important in mitochondrial function, and the ability of mitochondria to oxidize substrates after rotator cuff tear. While virtually no mitochondria were observed in the peripheral space of control muscles, extensive accumulation of peripheral mitochondria was noted after muscle injury (Figure 8A). There was no change in the ratio of mitochondrial DNA (mtDNA) to genomic DNA (gDNA, Figure 8B). Acylcarnitines and L-carnitine, which shuttle fatty acids between the cytosol and mitochondria, were reduced at all time points after injury (Figure 8C-F). Cardiolipins, which are phospholipids found almost exclusively within the inner mitochondrial membrane, were reduced 10 days after injury but recovered thereafter (Figure 8G). Glutathionylated proteins (Figure 8H), and two proteins that are critical in mitophagy, PINK1 (Figure 8I) and parkin (Figure 8J), were elevated after injury, along with ULK1 (Figure 8K) and p62 (Figure 8L) which play critical roles in mitophagy and autophagy in general. The hydrogen peroxide scavenging enzymes, peroxiredoxin (Prdx) 3 and peroxiredoxin 6, were elevated 10 days after injury, and for Prdx3 remained elevated at 30 and 60 days (Figure 8M-N). Succinate dehydrogenase (SDHA) and COX4 are two critical proteins in mitochondrial respiration, and while succinate dehydrogenase was reduced after injury, COX4 abundance was not different between control and injured muscles (Figure 8O-P). After assessing the abundance of proteins important in mitochondrial physiology, we then performed assays to determine the functional capacity of mitochondrial proteins to metabolize substrates. Complex I, II, and IV activity were reduced 10 days after injury, but recovered by 30 days (Figures 8Q-S). Similar results were observed when we evaluated the ability to oxidize pyruvate and palmitate, with an insufficiency 10 days after injury but recovery thereafter, with pyruvate oxidation nearly doubling 60 days after rotator cuff tear compared to control muscles (Figures 8T-U).

**Figure 8.**
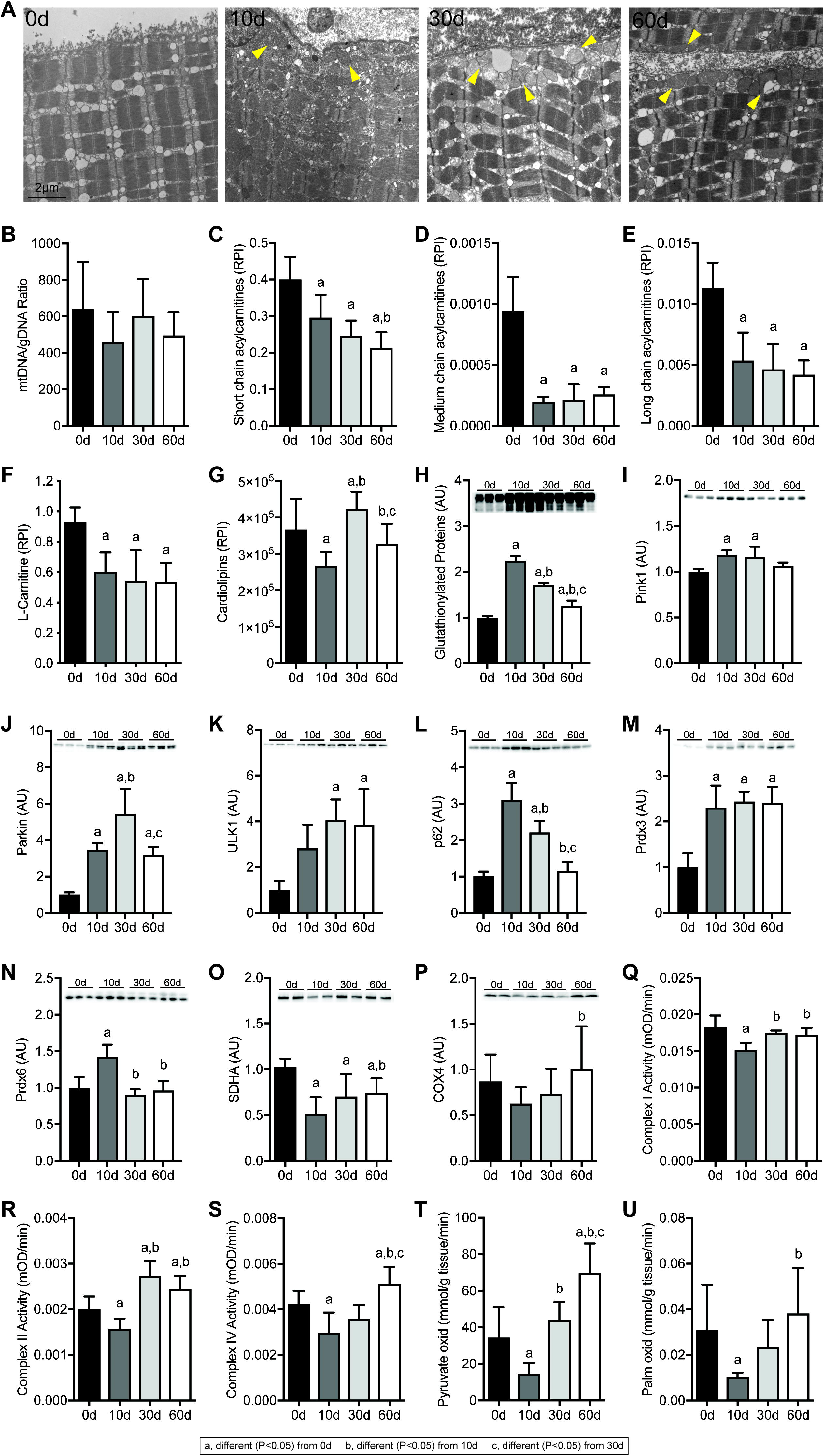
Changes in mitochondria abundance and function after rotator cuff tear. (A) Representative electron micrographs taken at the sarcolemma, with arrowheads demonstrating accumulation of peripheral, subsarcolemmal mitochondria. (B) Copies of mitochondrial DNA abundance to genomic DNA abundance. (C) Short chain (≤5 carbon) acylcarnitines, (D) Medium chain (6-12 carbon) acylcarnitines, (E) Long chain (13-20 carbon) acylcarnitines, (F) L-carnitine, and (G) Cardiolipins, as measured by mass spectrometry and presented as relative peak intensity (RPI). Quantification of Western blot band densitometry of (H) glutathionlylated proteins, (I) PINK1, (J) Parkin, (K) Ulk1, (L) p62, (M) Prdx3, (N) Prdx6, (O) SDHA and (P) COXIV protein levels, with representative blots shown as insets. Enzymatic activity of (Q) Complex I, (R) Complex II, and (S) Complex IV. Oxidation rates of (T) ^14^C-pyruvate and (U) ^14^C-palmitate. Data are presented mean+SD, N=6 muscles per group for B, N=5 muscles per group for C-F, N=10 muscles per group for G, N=3 muscles per group for H-P, N=4-6 muscles per group for Q-U. Post-hoc sorting (P<0.05): a, different from 0d; b, different from 10d; c, different from 30d.

## Discussion

Myosteatosis is a common pathological change observed in certain skeletal muscle groups following injury, and is particularly pronounced in the rotator cuff (7, 8). Fat accumulation is associated with greater muscle weakness and dysfunction in patients with rotator cuff tears (7, 11), and the mechanisms that lead to lipid accretion in myosteatosis are not well understood. Using a translational rat model of myosteatosis, we evaluated changes in muscle fiber force production and broadly profiled the changes in the muscle lipidome, metabolome, transcriptome and proteome, and utilized bioinformatics techniques to help identify potential factors that lead to fat accumulation in myosteatosis. Testing of muscle fiber force, along with electron micrographs, demonstrated a reduction in force production that accompanied disruptions to myofibril alignment and cytoskeletal architecture. Based on the transcriptional bioinformatics and supporting lipidomics, metabolomics, and electron micrographs, we then formulated the hypothesis that pathological lipid accumulation occurs in torn rotator cuff muscles due to mitochondrial dysfunction and reduced lipid oxidation. In support of this hypothesis, we observed a reduced capacity of mitochondria to oxidize lipids early in the injury process, along with transcriptional changes indicative of increased lipid droplet storage with reduced fatty acid uptake and mobilization from lipid droplet stores. Although mitochondrial function appears to recover at later time points, there is a general increase in glycolytic metabolites in muscles and a greater capacity to oxidize pyruvate. These findings support the notion that myosteatosis occurs due to a reduction in mitochondrial lipid oxidation, and not due to increased uptake of circulating lipids into the muscle. A summary of the pathological changes that occur after rotator cuff tear, and the relationships between these changes, is presented in Figure 9.

**Figure 9.**
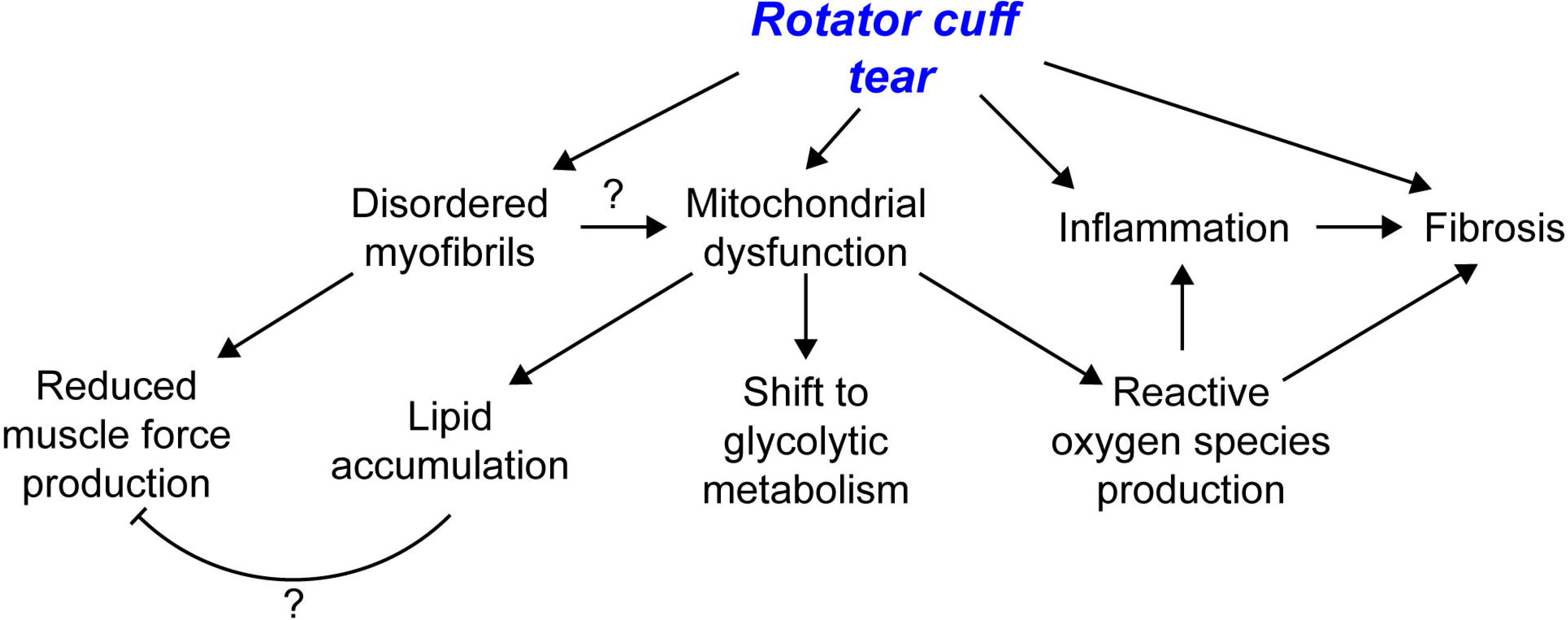
Overview of pathological changes after rotator cuff tear. An overview of the pathological changes observed and proposed that contribute to myosteatosis, weakness, and fibrosis after rotator cuff tear.

Shoulder weakness is a common observation in patients with rotator cuff tears, and this often does not recover even after successful repair of the torn tendon and completion of post-operative rehabilitation and strengthening programs (7, 11, 28). Satellite cells, which are myogenic progenitor cells that play an important role in regenerating muscle after injury, are found in equal abundance between untorn and fully torn rotator cuff muscles and display similar in vitro proliferative and fusion capacity (29), indicating that satellite cell dysfunction is unlikely to be responsible for the persistent dysfunction in patients who undergo rotator cuff repair. Despite apparently normal satellite cell activity, disordered sarcomere organization and reduced muscle force production has been reported in patients with chronic rotator cuff tears and animal models of chronic rotator cuff disease (1, 11, 12, 30, 31). However, less was known about subacute changes in contractile function after rotator cuff tear, and how this changes over time. In the current study, we observed a reduction in force production that occurred 10 days after the injury, continuing to decline by 30 days, with some recovery by 60 days. Electron micrographs demonstrated early sarcomere streaming, with the most grossly disorganized appearance of myofibrils at 30 days. By 60 days, myofibril organization improved, but with persistent z-disc malalignment that likely contributes to the reduced specific force at this time point. Numerous sarcomeric genes and proteins displayed marked changes in abundance at the 10 and 30 day time points, further suggesting that active myofibril remodeling occuring at these time points that tapers to some extent by 60 days. Genes and proteins involved with protein degradation, such as the E3-ligases MUSA-1, atrogin-1, MuRF-1, and NEDD4, and as the calcium-dependent protease m-calpain (32) were induced at 10 days. In agreement with these findings, the abundance of two important proteins in the 20S catalytic subunit of the 26S proteasome, PSMA6 and PSMB2 (33), were also increased after muscle injury. TGFβ, which can induce muscle atrophy by activating proteasomal protein degradation (34) was induced after injury, although surprisingly no differences in the closely related genes activin A and B, and a downregulation in myostatin (35), was observed. IL1β, which can promote inflammation and muscle atrophy through activating the NF_k_B pathway (36), was elevated across all injury time points.

IGF1, which is activated by exercise and can induce muscle hypertrophy through the Akt/p70S6K pathway (32) was upregulated after injury, but interestingly phosphorylation of the IGF1 receptor was reduced 30 and 60 days after rotator cuff tear. In type 2 diabetes, DG and ceramide-induced inflammation, lipotoxicity, and mitochondrial dysfunction are known to activate various serine/threonine kinases that block activation of the insulin receptor (37), which is closely related to the IGF1 receptor. Given the homologies between the two receptors, and the comparable lipotoxic and inflammatory environments, it is possible that similar mechanisms are inhibiting activation of the IGF1 receptor after rotator cuff tear. Phosphorylation of ERK1/2, which is a kinase downstream of IGF1 and several other mechanosensitive pathways in skeletal muscle (38), was also reduced after rotator cuff tear. IGF1 can both activate p70S6K pathway through an Akt/mTOR dependent phosphorylation of the T^389^ residue (39), and via ERK1/2 through the T^421^/S^424^ residues (40). Although p70S6K phosphorylation was reduced, activation of rpS6 which initiates translation (41) was increased after injury, suggesting an Akt/mTOR and ERK1/2 independent activation of translation in torn rotator cuff muscles. Although p70S6K is a well-known activator of rpS6, numerous other signaling pathways can also phosphorylate rpS6 (42), many of which are associated with cellular stress and inflammation, which is likely also occuring after rotator cuff tear. Although the mechanisms that determine which transcripts are selected for translation by ribosomes are not well understood, p70S6K activation is required for increasing muscle force during growth (43), suggesting a potential role for p70S6K in the translation of myofibrillar mRNAs. In the case of rotator cuff tears, since many patients continue to develop muscle atrophy and have persistent weakness after surgical repair and rehabilitation (7), it is possible that the reason the muscles of these patients fails to recover is due to suppression of the IGF1 signaling pathway that is normally activated in response to exercise.

In addition to muscle atrophy, fibrosis is a common feature of rotator cuff tears (44-46). In the current study, 10 days after injury there was an increase in the levels of the fibroblast marker FSP1, and several proteoglycans including lumican, osteoglycin, periostin, prolargin, thrombospondin 4, and vinculin. However, fibrillar collagens, measured by hydroxyproline levels, did not increase until 30 and 60 days after tear, corresponding to increases in the expression of several collagen transcripts and other ECM components at the same time points. These findings indicate that pathological changes to muscle fibers occur prior to substantial fibrotic changes in the ECM. As the accumulation of fibrotic ECM impairs lateral force transmission between muscle fibers which increases injury susceptibility during lengthening contractions (47), the changes that occur to the ECM likely further exacerbate weakness following rotator cuff tear.

While myosteatosis has been well documented in various types of muscle injuries and diseases, little is known about the ontogeny of fat accumulation in this condition (5, 6). In the current study, we identified a progressive increase in TG after injury, with TG levels at 60 days that were 3-fold higher than controls. This was consistent with observations of increased lipid accumulation observed in histology and electron microscopy. FFAs were also elevated at 30 and 60 days after rotator cuff tear. There was a downregulation in nearly all of the genes responsible for the transport of FFAs into the muscle cells, and the synthesis of TGs from FFAs (48), including CD36, ACSL1, ACSL6, AGPAT9/GPAT3, GPAM, AGPAT1, AGPAT3, AGPAT6, lipin 1-3, and DGAT1-2. Perilipin 2, which coats lipid droplets and appears to be important for lipid droplet growth and stability (49) was also upregulated after rotator cuff tear, while perilipin 5 which also coats lipid droplets but also helps to target fatty acids to mitochondria for oxidation (49), was downregulated. Consistent with these findings, genes involved with hydrolyzing triglyceride stored in lipid droplets into fatty acids (48, 50) were downregulated after injury, including ATGL and PNPLA3, hormone sensitive lipase, and monoglyceride lipase. These findings suggest that the intrafiber fat that accumulates in myosteatosis does not occur due to increased fatty acid transport into fibers or *de novo* synthesis of fatty acids from other substrates, but likely occurs due to a reduction in lipolysis.

In further support of this notion, mitochondrial lipid oxidation was also reduced 10 days after rotator cuff tear, with a corresponding reduction in the activity of complex I, II, and IV. It can be difficult to precisely measure mitochondrial abundance, especially since mitochondria often exist in large, interconnected networks throughout the muscle fiber (51), but mtDNA abundance and cardiolipin levels are often used to estimate mitochondrial mass (25, 52, 53). While we did not observe a difference in relative levels of mtDNA, a transient decrease in cardiolipins were observed 10 days after rotator cuff tear. Genes involved in the transport of fatty acids into the mitochondria and their conversion into fatty acyl-CoAs (ACSM5, ACSS1, CPT1b, carnitine acylcarnitine translocase, and CPT2), the β-oxidation of fatty acyl Co-As within the mitochondria (ACADS, ACADM, ACADL), and Kreb’s Cycle and oxidative phosphorylation genes (citrate synthase, aconitase, isocitrate dehydrogenase, 2-oxoglutarate dehydrogenase, succinyl CoA synthetase, SDHA, fumarase, malate dehydrogenase, NDUFS1, cytochrome C1, COX4, and ATP5G1) (48, 54, 55) were all downregulated after rotator cuff tear, with most remaining so through 60 days after injury. While *in vitro* assays demonstrated a reduction in fatty acid oxidation 10 days after injury, medium and long chain acylcarnitines, which serve as markers for the amount of fatty acids transported into mitochondria (55, 56), were reduced at all post-injury time points, suggesting a sustained reduction in fatty acid oxidation by mitochondria *in vivo*. Amino acids can be metabolized and eventually oxidized through the formation of short chain acylcarnitines (56), but these were also reduced after muscle injury, indicating that amino acid oxidation is not likely taking the place of lipid oxidation as a source of energy following muscle injury. Although mitochondrial lipid and protein oxidation appear to be reduced, ACOX1, which initiates the oxidation of fatty acids in peroxisomes and (57), and was also upregulated 10 days after rotator cuff tear. Peroxisomal oxidation of lipids also generates reactive oxygen species (57), which likely contribute to the inflammatory environment in muscles after injury.

Mitochondrial lipid oxidation appears reduced after rotator cuff tears, but metabolites involved in glycolysis are enriched 30 and 60 days after injury. Genes involved in glycolytic metabolism, such as phosphofructokinase, aldolase A, GAPDH, phosphoglycerate, and enolase 3 (58), were down at 10 days, but recovered to some extent by 60 days after injury, although glycolytic metabolites remained elevated. The net increase in glycolytic metabolites corresponded with a greater ability of mitochondria to oxidize pyruvate *in vitro*, and these findings in combination with the reduced abundance of acylcarnitines suggest muscle fibers are using much less lipid for oxidation. Further, while there was an accumulation in glycolytic metabolites, no apparent change in myosin isoform expression is present as would be expected with a major fiber type transition occurred. The overall lipid accumulation and shift to a glycolytic phenotype that we observed in this study is similar to the findings in a mouse model of skeletal muscle specific CPT1b deletion, in which mitochondria had a drastically reduced ability to take up long chain fatty acids, resulting in an accumulation of cytosolic lipid droplets, as well as elevated levels of TG and ceramides, and an increase in glycolytic metabolism (59).

Several reports involving animal and human models of skeletal muscle unloading and denervation have identified profound mitochondrial dysfunction and accompanying increases in ROS production that occur subsequent to the injury (60-63). We also observed increased markers of ROS in injured muscles, including elevated levels of glutathionylated proteins, as well as an increase in the abundance of Prdx3 and Prdx6 which scavenge hydrogen peroxide in the mitochondria and cytoplasm, respectively (64, 65). The production of ROS by dysfunctional mitochondria, as suggested by the elevation in Prdx3, likely further exacerbates the inflammatory environment within injured rotator cuff muscle fibers, as elevated ROS has been linked to inhibition of protein synthesis signaling pathways, the induction of proteolytic and autophagic pathways, and fibrosis (66, 67). To address the dysfunctional mitochondria, injured rotator cuff muscle fibers do appear to be activating mitophagy, as evidenced by increased levels of PINK1 and Parkin which target depolarized mitochondria for breakdown in autophagosomes, in a processed mediated by ULK1 (68). p62, which plays an important regulatory role in both mitophagy and general autophagy (35), was also markedly elevated at all time points after rotator cuff tear. However, although there appears to be an increase in mitophagy, key genes involved in mitochondrial biogenesis, including PGCla, NRF1, and TFAM (69) were downregulated after injury, as were the mitochondrial fusion genes Mfn1 and Mfn2. We also noted an accumulation of peripheral segment mitochondria in torn rotator cuff muscles.

In the classical two mitochondrial population model, a collection of peripheral mitochondria have decreased oxidative capacity compared to the physically distinct pool of mitochondria in the myofibrillar space (70-72). In the newly emerging view of mitochondria, where mitochondria are thought to exist in a dynamic reticulum extending from the sarcolemma to the myofibrils, and actively undergoing fusion and fission in continuous networks throughout the cell, peripheral segment mitochondria have a greater abundance of proton motive proteins, whereas the connected intermyofibrillar mitochondria located in the I-band have more ATP generating proteins for use by sarcomeres (51, 73). In the current study, the abnormal spherical mitochondria in the peripheral space of injured muscles at 30 and 60 days after rotator cuff tear could reflect a mitochondrial network that has been mechanically and biochemically disrupted, and this is supported from observations in yeast in which the small mitochondria that bud off of larger networks of mitochondria have a spherical appearance (74). While the nature of mitochondrial dysfunction in torn rotator cuff muscles remains unknown, as the continuum of mitochondrial networks that normally exist in healthy skeletal muscle likely requires extensive support from the cytoskeleton of the fiber to maintain shape, we posit that the highly disrupted myofibril architecture that occurs as a result of rotator cuff tear likely interferes with the ability to form a stable intermyofibrillar network, resulting in mitochondrial dysfunction and subsequent lipid accumulation.

There are several limitations to this study. While the rat model is commonly used in the study of rotator cuff pathology, the interfiber fat accumulation phenotype is less pronounced than what is observed in patients with chronic rotator cuff tears. Additionally, humans and large animals of rotator cuff injury develop a combination of inter- and intrafiber lipid accumulation, with the intrafiber accumulation occuring through an expansion of adipocytes. This rat model has virtually no adipocytes that accumulate after injury, which limits our analysis to changes in fat accumulation that occur within muscle fibers. We only evaluated rats with a rotator cuff tear, and did not evaluate changes that occur within the muscle after the tear is repaired. Three postinjury time points were selected to look at subacute, early chronic, and chronic changes in the muscle, but extending the evaluation to early and later time points is likely to add additional information about the pathology. Finally, while only male rats were studied, but we think the results are informative about rotator cuff pathology in both sexes.

Rotator cuff tears are among the most prevalent upper extremity disorders, and can result in profound pain and disability that persist despite surgical repair (7, 9, 75). While surgical techniques have evolved to improve the repair of the torn tendon back to its insertion on the humerus, our ability to treat the extensive muscle atrophy and myosteatosis that occur subsequent to the tear is limited (1, 8). The findings from this study demonstrate that the fat which accumulates in torn rotator cuff muscles likely occurs due to deficits in mitochondrial lipid oxidation, and that the accumulation of fats likely leads to a lipid-induced pro-inflammatory and lipotoxic state which reduces regeneration. Further exploration of therapies to enhance mitochondrial function and increase lipid oxidation may lead to improvements for patients with rotator cuff tears and other myosteatosis-related conditions.

## Acknowledgements

This work was supported by grants F31-AR065931, R01-AR063649, R01-DK107397, R03-DK109888, and U24-DK097153 from the National Institutes of Health. We also acknowledge technical contributions from Dr. Richard McEachin, Dr. James Markworth, and Mr. Jacob Swanson.

## Author Contributions

JPG, KF, BM, CLM designed research; JPG, AHQ, PJF, BM, CLM performed research; ANM contributed analytic tools; JPG, AHQ, PJF, KF, BM, CLM analyzed data; JPG and CLM wrote the paper.

## Non-Standard Abbreviations

2PG+3PG: 2-phosphoglycerate and 3-phosphoglycerate
ADP: adenosine diphosphate
AMP: adenosine monophosphate
ATP: adenosine triphosphate
CIT+ICIT: citrate and isocitrate
CMP: cytidine monophosphate
F6P+G6P: fructose-6-phosphate and glucose-6-phosphate
FAD: flavin adenine dinucleotide
GDP: guanosine diphosphate
GMP: guanosine monophosphate
NAD: nicotinamide adenine dinucleotide
NADP: nicotinamide adenine dinucleotide phosphate
R5P+X5P: ribulose 5-phosphate and xylulose 5-phosphate
UDP: uridine diphosphate
UMP: uridine monophosphate
UTP: uridine triphosphate

